# Data-driven multiplexed microtomography of endogenous subcellular dynamics

**DOI:** 10.1101/2020.09.16.300392

**Authors:** YoungJu Jo, Hyungjoo Cho, Wei Sun Park, Geon Kim, Donghun Ryu, Young Seo Kim, Moosung Lee, Hosung Joo, HangHun Jo, Sumin Lee, Hyun-seok Min, Won Do Heo, YongKeun Park

**Affiliations:** Department of Physics, Korea Advanced Institute of Science and Technology (KAIST), Daejeon 34141, Republic of Korea; KAIST Institute for Health Science and Technology, KAIST, Daejeon 34141, Republic of Korea; Tomocube Inc., Daejeon 34051, Republic of Korea; Department of Biological Sciences, KAIST, Daejeon 34141, Republic of Korea; Center for Cognition and Sociality, Institute for Basic Science (IBS), Daejeon 34141, Republic of Korea; Present address: Departments of Applied Physics and of Biology, Stanford University, Stanford, CA 94305, USA

## Abstract

Simultaneous imaging of various facets of intact biological systems across multiple spatiotemporal scales would be an invaluable tool in biomedicine. However, conventional imaging modalities have stark tradeoffs precluding the fulfilment of all functional requirements. Here we propose the refractive index (RI), an intrinsic quantity governing light-matter interaction, as a means for such measurement. We show that major endogenous subcellular structures, which are conventionally accessed via exogenous fluorescence labeling, are encoded in 3D RI tomograms. We decode this information in a data-driven manner, thereby achieving multiplexed microtomography. This approach inherits the advantages of both high-specificity fluorescence imaging and label-free RI imaging. The performance, reliability, and scalability of this technology have been extensively characterized, and its application within single-cell profiling at unprecedented scales has been demonstrated.

## Introduction

Imaging is the process of mapping a variable, called *contrast*, in space and time. The tradeoffs between different contrast mechanisms fundamentally determine the distinct characteristics of each imaging modality (Mertz, 2019). In biomedicine, fluorescence (FL) has been a canonical imaging contrast over several decades for visualizing specific elements within biological systems, powered by chemical, immunological, and genetic labeling strategies (Lichtman and Conchello, 2005). Despite the excellent biochemical specificity, however, a number of drawbacks are associated with FL: photobleaching and phototoxicity (limited temporal window), spectral overlap (limited multiplexing), variability of labeling quality (limited reproducibility and potential bias), and exogenous labeling-induced side effects (potential perturbation of endogenous biology).

Recent advances in machine learning triggered an interesting approach, known as cross-modality inference or *in silico* labeling. This approach achieves FL contrast by measuring another contrast with complementary characteristics (Christiansen et al., 2018; Nygate et al., 2020; Ounkomol et al., 2018; Rivenson et al., 2019). Most notably, the paring of images from simultaneous FL and bright-field (BF) microscopy, based on light absorption contrast, was used to train neural networks to convert BF images into FL images (Christiansen et al., 2018; Ounkomol et al., 2018) (“BF2FL”). Despite the ingenuity of this method enabling computational staining of unlabeled samples, the *minimal absorption* at the cellular and subcellular levels raises a fundamental question about BF2FL: is light absorption an optimal contrast mechanism for cross-modality inference?

An alternative approach utilizes optical phase delay as the measured contrast (Nygate et al., 2020; Rivenson et al., 2019). Unlike absorption, phase distributions have a significant contrast in space even at the subcellular level, and forms the basis of phase-contrast microscopy. Emerging quantitative phase imaging (QPI) technologies measure the phase images of unlabeled samples with high sensitivity (Park et al., 2018), which could be paired with FL images for cross-modality inference. Although the improved contrast mechanism is expected to facilitate robust performance, so far, this method has not been extended beyond 2D imaging, which has significant limitations for the detailed characterization of subcellular dynamics.

In this study, we report a new technology for data-driven multiplexed microtomography of endogenous subcellular structures and dynamics across various spatiotemporal scales. We fundamentally improved the cross-modality inference framework by introducing the contrast mechanism of the 3D refractive index (RI), which is an endogenous quantity governing light-matter interaction including both absorption and phase delay. The diffraction-limited 3D RI tomograms, measured by high-speed multi-angle holography (or holotomography), enabled the scalable inference of multiple 3D FL tomograms for the corresponding subcellular targets (“RI2FL”). Importantly, we found that RI2FL generalizes across cell types, outperforming BF2FL in all quantitative measures. Moreover, Bayesian uncertainty quantification schemes provided measure of spatiotemporal reliability. We demonstrated a proof-of-concept application of RI2FL in cell biology and high-throughput screening via unprecedented single-cell profiling.

## Results

We sought to determine the quantitative relations linking RI distribution to subcellular targets in a data-driven manner by training 3D convolutional neural networks to translate a RI tomogram into FL tomograms corresponding to multiple subcellular targets (Figure 1A). We first created a large-scale dataset consisting of ~1,600 3D RI tomograms (at 532 nm wavelength) and the corresponding 3D FL tomograms from 6 subcellular targets (actin, mitochondria, lipid droplets, plasma membranes, nuclei, and nucleoli) and 6 eukaryotic cell types (NIH3T3, COS-7, HEK 293, HeLa, MDA-MB-231, and astrocyte) using standardized holotomographic microscopes equipped with FL channels (Kim et al., 2017) (Methods). To train the networks, we used a subset of NIH3T3 tomograms only and held out all other data, in order to test the generalization of the discovered RI-target relations across cell types (Table S1).

**Figure 1.**
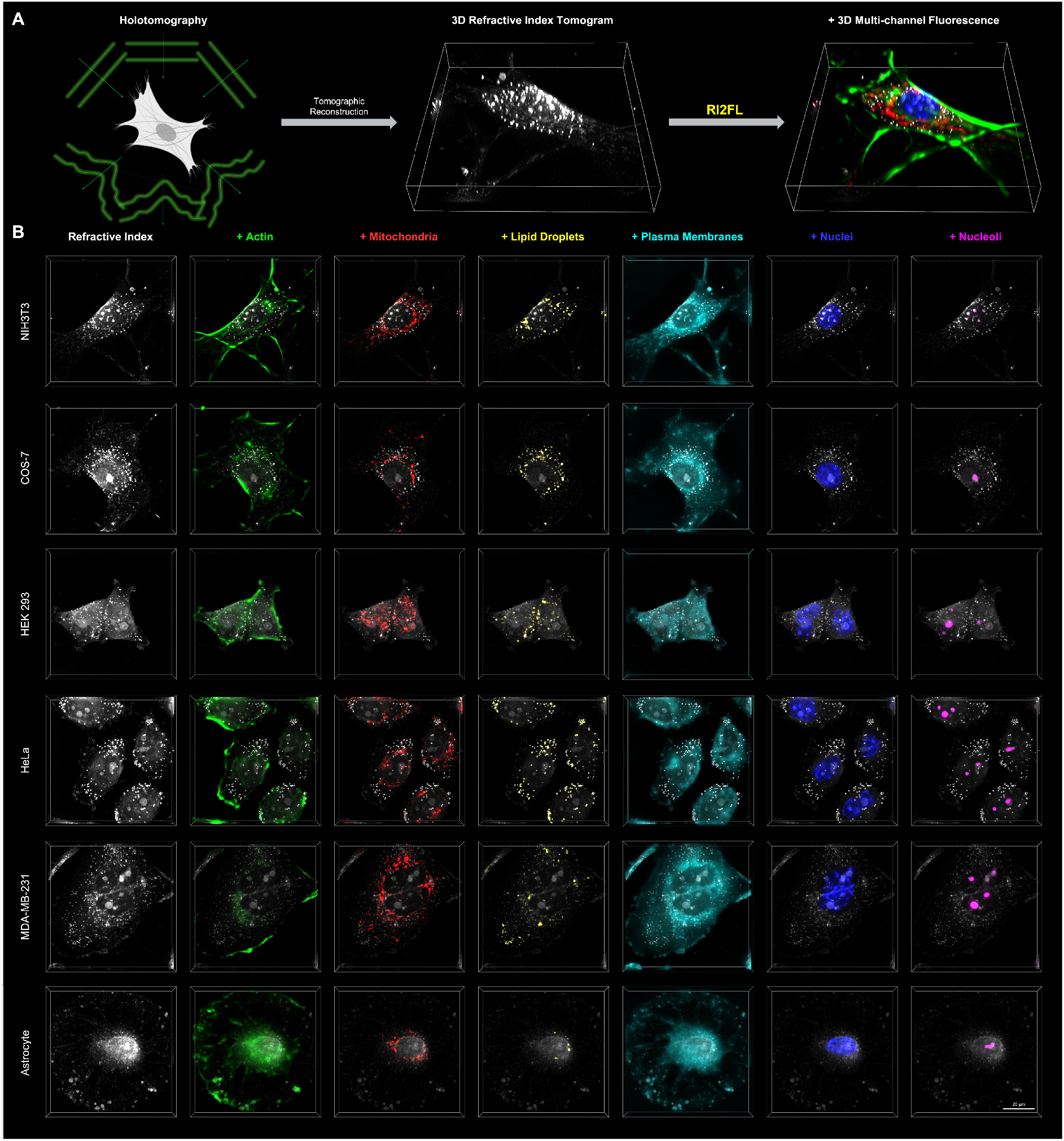
Data-driven microtomography of multiple subcellular structures. (A) Concept of RI2FL. (B) Examples of RI2FL across cell types and subcellular targets. All bounding boxes represent a volume of 76.8 × 76.8 × 12.8 μm^3^.

The distinct nature of RI and FL presents several experimental and computational challenges. Firstly, while RI is an absolute and unbiased quantity independent of the experimenter or instrument, FL signals are heavily dependent on labeling quality, illumination power, and exposure time (Lichtman and Conchello, 2005; Mertz, 2019). To amend this, we implemented tight quality control procedures carried out by trained cell biologists throughout the data acquisition and processing pipeline, thereby establishing ground truth subcellular targets defined by 3D FL (Methods). Secondly, the drastic differences between the FL channels require the painstaking optimization of individual target-specific network architectures. We instead utilized a single, highly flexible network architecture for all subcellular targets, powered by a large-scale neural architecture search, which was also potentially advantageous for an extension to additional subcellular targets. (Figure S1). Thirdly, high-resolution 3D RI tomograms have enormous memory demands infeasible for most GPUs. To avoid this, we assumed that the target-specific patterns could be identified from local (~10 μm) distribution of RI, and implemented patch-based parallel processing (Figure S2). With these strategies in hand, we successfully trained the networks for RI2FL inference (Figure 1B, Figure S3, and Video S1).

We characterized the performance of RI2FL by quantitatively comparing the inferred and ground truth FL in the held-out dataset. The prediction accuracy across cell types is illustrated in Figure 2A. Strikingly, not only NIH3T3 cells but also all other cell types, which were never presented to the networks during training, showed high performance. In particular, an excellent accuracy for astrocytes, which were obtained from primary cultures unlike other immortalized cell lines, strongly supported that RI2FL captured fundamental RI-target relations generalizing across cell types. The per-target performance is presented in Figure 2B. While the high accuracy for nuclei and lipid droplets is consistent with the high RI contrast of these targets (Lee et al., 2019; Park et al., 2020), all the remaining targets, which are hardly recognizable via the visual inspection of RI tomograms, showed comparable performances. In addition, in order to rule out the labeling-induced bias of the dataset (which is unlikely due to the low density of fluorophores (Yoon et al., 2018)), we performed RI2FL with unstained samples, and then stained the samples to obtain the corresponding FL ground truth, which showed consistent results (Figure S4). Together, our results confirmed the seamless identification of endogenous subcellular targets by RI2FL (see Table S2 for additional performance metrics).

**Figure 2.**
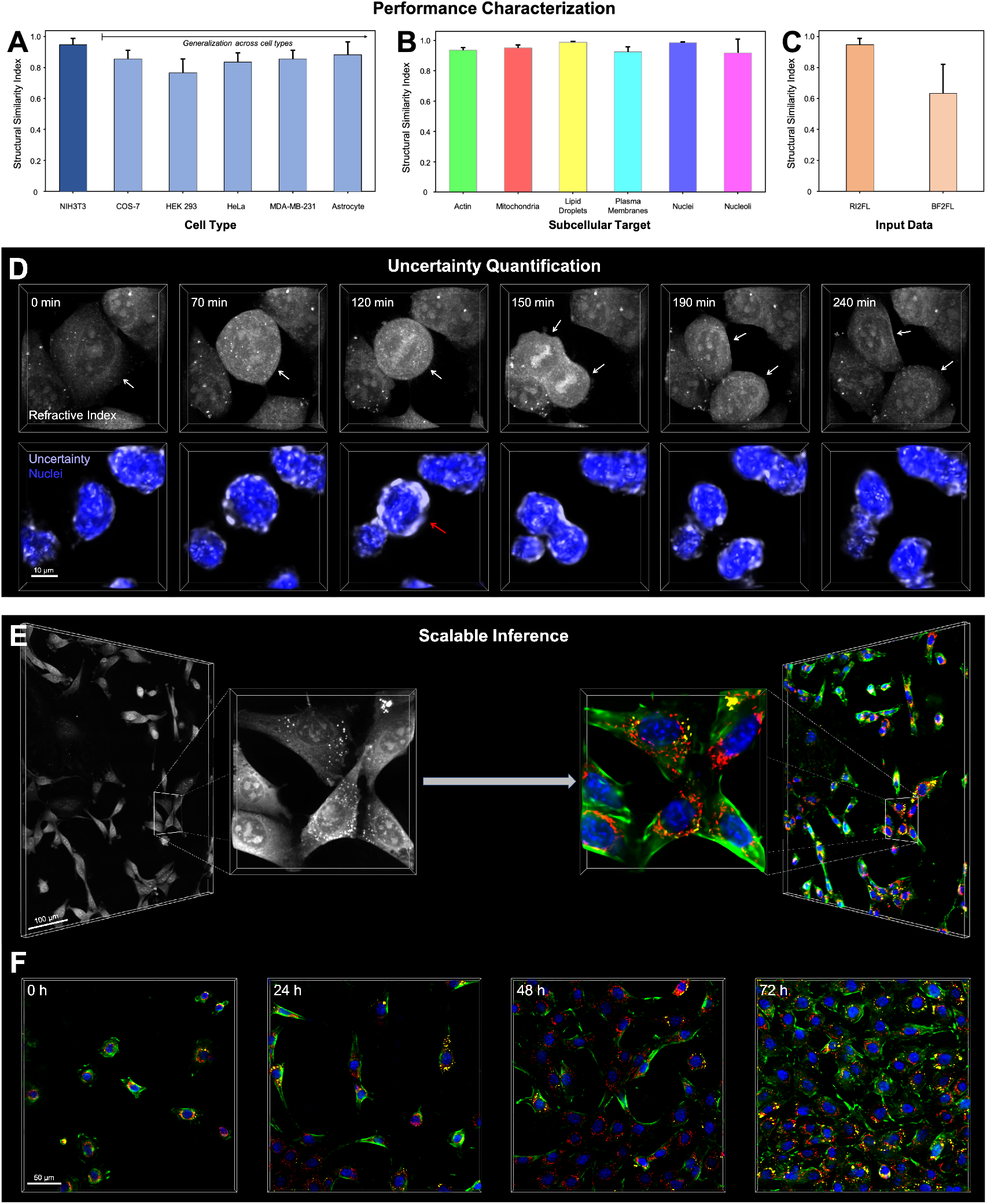
Performance, reliability, and scalability. (A-C) RI2FL performance quantification across (A) cell types, (B) subcellular targets, and (C) methods. All quantifications were conducted on the held-out data. Mean±S.D.; further statistics are provided in Table S2. (D) Uncertainty quantification example. Top: The dynamics of cell division observed with timelapse RI tomograms. White arrows indicate the dividing cell and its daughter cells. Bottom: Corresponding nuclei inference and its data uncertainty maps. The red arrow highlights the larger uncertainty due to the putative breakdown of the nuclear envelope, which is specific to this cell state. The bounding boxes represent a volume of 49.2 × 49.2 × 12.8 μm^3^. (E) Scaling up RI2FL to large FOV tomograms exploiting the shift-invariance of the RI-target relations. The larger bounding boxes represent a volume of 480 × 480 × 13 μm^3^. (F) Scaling up RI2FL in space and time, further exploiting the time invariance of the RI-target relations. The bounding boxes represent a volume of 307×307×13 μm^3^. In (E and F), only four FL (actin, mitochondria, nuclei, and lipid droplets) channels are shown for visual clarity.

Next we quantified the advantage of RI over BF (Christiansen et al., 2018; Ounkomol et al., 2018). Because RI includes both absorption and phase delay information, one can reconstruct stacked BF intensity images from RI tomograms using Fourier optics (Mertz, 2019) (Methods). We trained BF2FL networks with the reconstructed BF images and the corresponding FL tomograms (Figure S3), and compared the performances of BF2FL and RI2FL (Figure 2C). Clearly, there was a considerable margin between the two methods. This is not surprising because the optical information in stacked BF images is only a tiny subset of the full RI information, particularly in cellular systems where phase delay dominates over absorption (Park et al., 2018).

Despite the exciting opportunities provided by RI2FL in addition to previous cross-modality inference approaches, the extent to which we can “trust” the model predictions in space and time has not been clear. We aimed to elucidate this by applying the recent advances in Bayesian deep learning (Gal and Ghahramani, 2016; Wang et al., 2019). To this end, we quantified the uncertainty maps to guide the endusers with “error bars” accompanying the FL predictions. Intuitively, uncertainty can be estimated as the voxel-wise variability of predictions upon perturbation of the data or model (Figure S5). An example demonstrating uncertainty quantification in RI2FL is presented in Figure 2D (also see Video S2). During animal cell division, the nuclear envelope breaks down to facilitate the separation of aligned chromosomes by the spindle apparatus. This specific event, which is rare in the training dataset, makes nuclei prediction by RI2FL particularly challenging around the periphery of the nuclei. The specific increase in uncertainty at this stage (red arrow) casts a cautionary signal for downstream analyses. As such, uncertainty quantification can provide spatiotemporal reliability measures for the end-users, on top of the holistic accuracy metrics. The uncertainty maps can also guide data collection to strengthen the model.

RI2FL is intrinsically scalable in space and time because the RI-target relations are shift- and time-invariant. The trained models, together with the patch-based processing (Figure S2), can be readily applied to the large field-of-view (FOV) RI tomograms obtained by image stitching or high space-bandwidth product techniques (Baek et al., 2019; Zheng et al., 2013). We successfully operated RI2FL for tomograms with large FOVs up to 480×480×13 μm^3^ without trading off the high spatial resolution (Figure 2E). In addition, RI2FL can be sequentially applied to time-lapse large-FOV tomograms, as we demonstrated in the recordings up to 72 hours (Figure 2F and Video S3). Importantly, there is no theoretical upper limit in the spatial and temporal scales by virtue of holotomography that is free from photobleaching and phototoxicity; the only practical limitations are computing time and memory, which linearly scale with the data dimensions.

One exciting application of RI2FL is time-resolved hybrid single-cell profiling for use in cell biology and high-throughput screening. The image-based profiling of single cells with standardized data acquisition and interpretable feature extraction has provided insights into new phenotypes and cellular heterogeneity, thereby complementing genomics (e.g., Cell Painting with CellProfiler (Bray et al., 2016)). While this capability is critically dependent on highly multiplexed FL imaging, the spectral overlap issue limits such measurement to *fixed* cells via multi-round imaging. On the other hand, multiplexed microtomography with RI2FL in intact *living* cells allows for the time-resolved profiling of single-cell phenotypes. Furthermore, RI contributes additional information orthogonal to FL. Traditionally, RI has been a uniquely suitable modality for ultrasensitive quantification of subcellular mass (Barer, 1953), which is particularly relevant for the study of cell cycle and growth (Cooper et al., 2013; Mir et al., 2011). This quantitative nature of RI can be synergistically combined with the specificity of FL to access new dimensions for single-cell profiling (Figure 3A; Methods).

**Figure 3.**
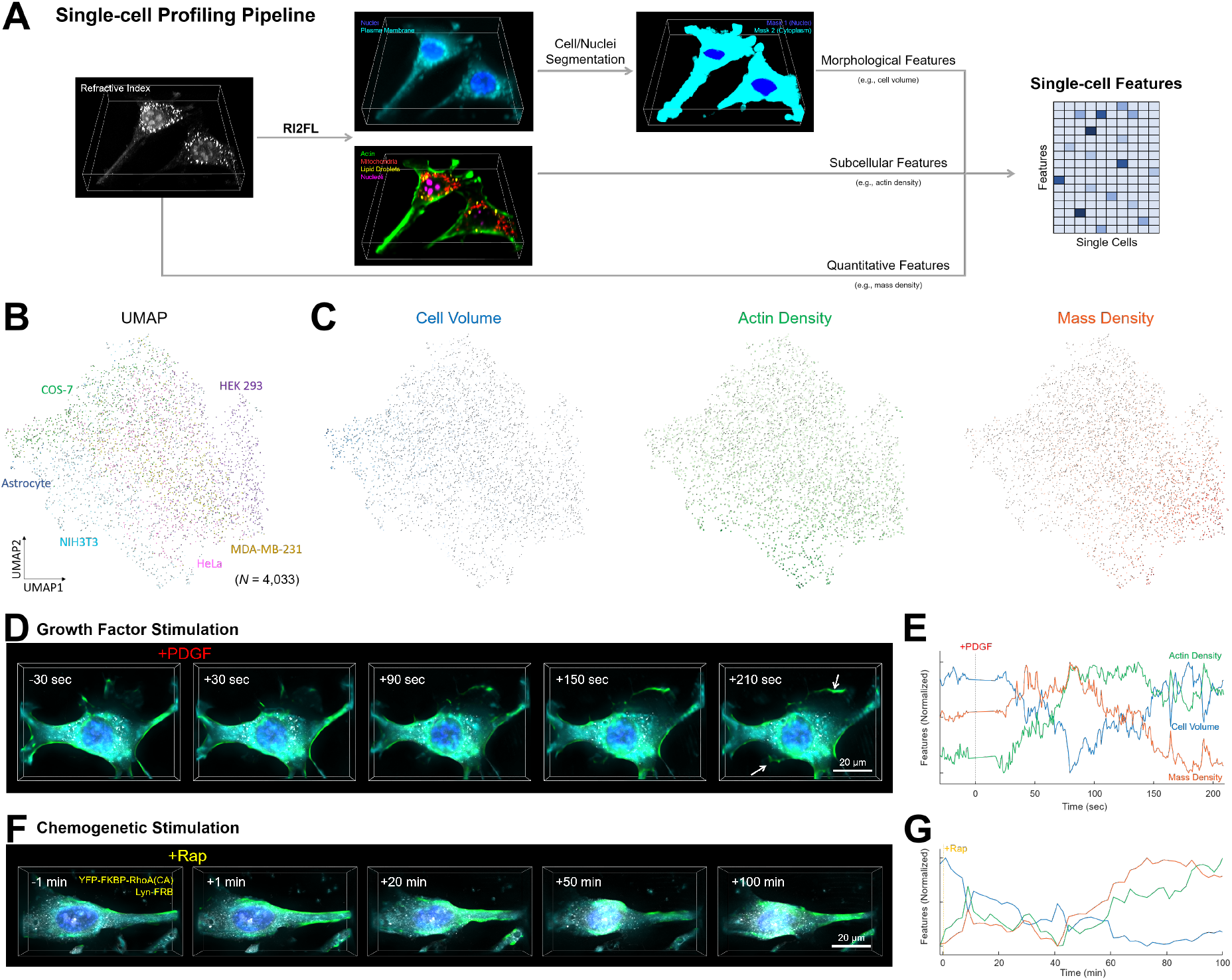
Application to single-cell profiling at unprecedented spatiotemporal scales. (A) Fully automated single-cell profiling pipeline based on RI2FL. The bounding boxes represent a volume of 76.8 × 76.8 × 12.8 μm^3^. (B) Unsupervised embedding of single cells based on the extracted single-cell features. (C) Visualization of three representative features mapped onto single cells. (D–G) Perturbation experiments with (D and E) growth factor and (F and G) chemogenetic stimulation. (D and F) Time-lapse RI2FL. RI and three FL (actin, nuclei, and plasma membranes) channels are shown. The bounding boxes represent a volume of (D) 76.8 × 55.4 × 12.8 μm^3^ and (F) 76.8 × 76.8 × 12.8 μm^3^. (E and G) The corresponding feature dynamics over time. Only three out of 65 features are shown for visual clarity.

For a sanity check, we profiled ~4,000 single cells detected in the dataset with the six cell types. Our fully automated pipeline robustly segmented individual cells and extracted a variety of interpretable features defined by morphology, FL, RI, or their combinations (Table S3). We defined a minimally redundant set of 65 features spanning the representative facets of the single cells, while one can readily define thousands of single-cell features with the current level of multiplexing (Bray et al., 2016). The unsupervised low-dimensional embedding of the single-cell features revealed an intriguing variability across and within the cell types (McInnes et al., 2018) (Figure 3B). Three exemplary features underlying this variability are shown in Figure 3C. Astrocytes generally had large cell volumes, consistent with their complex stellar morphology. NIH3T3 cells, which are fibroblasts with characteristic actin structures, showed a high actin density. HEK 293 cells had a high mass density, which can be attributed to the high rates of protein production by this cell type. Notably, for these three, as well as all the other features, an enormous variability was observed within each cell type, which is partially dependent on the cell cycle, making single-cell profiling attractive.

Next, we proceeded to time-resolved interrogations. For proof-of-concept, we carried out a series of perturbation experiments. Firstly, NIH3T3 fibroblasts were stimulated with platelet-derived growth factor (PDGF) to promote cell growth in a physiological manner (Martin et al., 2014), while observing the cells with a high volume rate (0.8 sec/volume) exploiting the high speed of holotomography without photobleaching and phototoxicity. The inferred FL channels clearly visualized lamellipodia formation (white arrows) and actin reorganization in 3D at a time scale of minutes (Figure 3D and Video S4). The time-resolved profiling was able to measure the fast dynamics of the features in response to PDGF stimulation (Figure 3E). This measurement represents a new regime in cell biology that has been previously inaccessible due to technical limitations. The temporal resolution can be readily improved beyond the video rate (Kim et al., 2013).

To determine the roles of the distinct signaling pathways, we specifically targeted RhoA, a Rho family small GTPase downstream of PDGF, using chemogenetics (Inoue et al., 2005). The rapamycin-induced formation of the FKBP-FRB-rapamycin complex recruited constitutively active RhoA to the plasma membrane (Figure 3F and Video S5). Unlike PDGF, RhoA stimulation specifically promoted the formation of actin stress fibers, resulting in characteristic cell morphology. Consistently, only a subset of the features illustrated in Figure 3E showed similar dynamics to PDGF stimulation (Figure 3G). The whole variety of features, including those based on multiple FL channels, may be relevant for studying inter-organelle interactions during these cellular responses. Taken together, our results demonstrate that RI2FL provides a powerful means by which to quantitatively profile single living cells via simultaneous access to a variety of information.

## Discussion

In summary, we developed and extensively characterized RI2FL, a scalable framework with which to infer endogenous subcellular structures and dynamics from 3D RI tomograms. The high performance of this approach was found to result from the nature of RI, which encompasses both absorption and phase delay information. Together with the uncertainty quantification schemes to measure the prediction reliability in space and time, RI2FL represents a powerful platform technology for cell biology and high-throughput screening, as demonstrated by its capacity for time-resolved single-cell profiling.

The high-dimensional observation and perturbation of single-cell dynamics at scale would facilitate the systems-level understanding of cellular behavior and decision-making. So far, most studies in systems biology have been relying on snapshot (e.g., transcriptomics or multi-round imaging) or low-dimensional time-series (e.g., one or two fluorescence biosensors) measurement of cellular states, which had significant limitations to infer the network-based logic governing cellular dynamics. With the highdimensional time-series measurement (or single-cell state space trajectories) in hand, one could directly infer the underlying dynamical systems at the single-cell level, analogously to systems neuroscience (Pandarinath et al., 2018). We are currently applying this approach to search for the dynamical phenotypes specific to clinically-relevant cellular malfunctions (e.g., cancer) or to high-efficacy drug actions (Topol, 2019).

We believe that this study, at least partly, addresses a long-sought goal of the label-free imaging community: biochemical specificity by RI. While the overlapping RI values of many subcellular structures have precluded RI-based imaging beyond nuclei and lipid droplets (Figure S6), we previously proposed that the *spatial distribution* of RI may encode enough information to infer these structures (Jo et al., 2019), as experimentally demonstrated in the present study. We look forward to extending this technology to tissues and ultimately to *in vivo* applications, synergizing with new approaches to 3D QPI in highly scattering systems (Chen et al., 2020; Lim et al., 2019).

An important next step would be reverse-engineering the trained models to interpret the discovered RI-target relations (Doshi-Velez and Kim, 2017; Sussillo and Barak, 2013) (Figure S7). At the moment, the feasibility of RI2FL for a new target can be tested only empirically by target-by-target training and characterization. The interpretability might reveal general principles governing light-matter interaction in biological systems and clarify the fundamental limits of RI2FL as well as other cross-modality approaches.

## Acknowledgments

We thank the members of KAIST Biomedical Optics Laboratory for helpful discussions. This work was supported by KAIST, BK21+ program, Tomocube Inc., and National Research Foundation of Korea (2017M3C1A3013923, 2015R1A3A2066550, and 2018K000396). Y.-J.J. was supported by a KAIST Presidential Fellowship, an Asan Foundation Biomedical Science Scholarship, and a Stanford Bio-X Bowes Fellowship.

## Author Contributions

Y.-J.J. and Y.-K.P. conceived the idea. Y.-J.J. coordinated the project and carried out all analyses. Y.-J.J., H.C., and H.-S.M. designed the deep learning pipeline, and H.C. implemented and optimized the pipeline. Y.-J.J., W.S.P., and W.D.H. designed the main experiments, and W.S.P. collected the data. W.S.P. and W.D.H. established the molecular biology and imaging protocols. G.K., D.R., Y.S.K., and M.L. contributed to processing the data. H.J., H.-H.J., and S.L. collected and processed the large FOV data. Y.-J.J., H.C., W.S.P., and Y.-K.P. wrote the manuscript with input from all co-authors. Y.-K.P. supervised all aspects of the work.

## Methods

### Sample preparation

NIH3T3 (ATCC CRL-1658), COS-7 (ATCC CRL-1651), HEK 293 (ATCC CRL-1573), and HeLa (ATCC CCL-2) cells, as well as primary-cultured astrocytes, were maintained in Dulbecco’s modified Eagle’s medium (DMEM; ATCC 30-2002) supplemented with 10% fetal bovine serum (Life Technologies) and 100 U/mL penicillin-streptomycin at 37°C in a 5% CO_2_ incubator. For MDA-MB-231 (ATCC HTB-26) cells, DMEM was replaced with DMEM/F12 medium (Gibco). Astrocytes were obtained by cortical and hippocampal dissection of embryos from C57BL/6J mice (Jackson Laboratory), followed by astrocyte enrichment with a glial culture medium. All animal procedures were performed according to the guidelines of the Animal Care and Use Committee at KAIST.

The fluorescence labeling strategies were as follows. Actin was stained by phalloidin (A12379; Invitrogen) or genetically labeled by expressing mCherry-Lifeact (constructed by inserting the F-actin peptide-encoding sequence into the mCherry-C1 vector). Mitochondria were stained by MitoTracker Red CMXRos (M7512; Invitrogen). Lipid droplets were stained by LipiDye (FDV-0010; Funakoshi). Plasma membranes were stained by CellMask (C10046; Invitrogen) or genetically labeled by expressing GFP-MEM. Nuclei were stained by Hoechst 33342 (H3570; Life Technologies). Nuclei were genetically labeled by expressing FBL-mCherry. For the chemogenetic stimulation experiment, Lyn-FRB and YFP-FKBP-RhoA(CA) were co-expressed in NIH3T3 cells. All genetic labeling processes were driven by CMV promoters. The cells were transfected via electroporation (Neon Transfection System; Invitrogen) under the following conditions: voltage, 1,280 V; pulse width, 20 ms; number of pulses, 2.

Holotomography-optimized cell culture dishes (Tomodish; Tomocube Inc.) were seeded with approximately 450,000 cells per dish. The culture dishes were coated with 0.01% poly-D-lysine for 15 minutes, washed three times with distilled water, and fully dried before usage.

### Imaging and perturbation

Holotomography was conducted using standardized microscopes (HT-2; Tomocube Inc.) implementing optical diffraction tomography (ODT) to solve the RI-thickness coupling problem in phase images. The principles and implementations of ODT have been extensively reviewed elsewhere (Park et al., 2018). The specific implementation used here was based on holographic field retrieval with multi-angle Mach-Zehnder interferometry using coherent 532 nm laser light steered by a digital micromirror device. The 3D RI tomograms were reconstructed by mapping the measured field information to the 3D Fourier space and filling the missing cone using the non-negativity-constrained iterative algorithm. The acquisition time for a single volume was less than a second. In addition, we optionally utilized the three FL channels to measure the ground truth 3D FL tomograms using stacked widefield acquisition with a ~0.3 μm step size followed by 3D deconvolution (Kim et al., 2017) (excitation center wavelengths: 385 nm, 470 nm, and 565 nm). Excitation light intensity and exposure time were manually adjusted by trained cell biologists to clearly visualize the target structures. For the perturbation experiments, the concentrations of PDGF (PDGF-BB, PeproTech) and rapamycin (Calbiocam) in the imaging medium were 10 nM and 0.5 μM, respectively.

### Data processing

All tomograms were resized to have a voxel size of 0.15×0.15×0.2 μm^3^ before inference. The default FOV used in training and evaluation was 512×512×64 voxels, which corresponds to a volume of 76.8×76.8×12.8 μm^3^ volume. This volume was then further subdivided for patched-based processing. The large FOV RI tomograms were obtained by offline 3D stitching after the rapid acquisition of slightly overlapping tiles of FOVs. Specifically, we estimated the tile-to-tile displacement using the phase correlation of the single-FOV RI tomograms, and the overlapping volumes were processed by image blending. The RI values were clipped into the range between 1.337 and 1.390. While we targeted the identical FOV for the RI and FL tomograms by sharing most optical path, we noticed a small axial discrepancy due to the intrinsic differences between the modalities (e.g., aberration and latency). In order to secure voxel-wise correspondence to facilitate supervised learning and evaluation, we estimated and corrected this discrepancy using the axial cross-correlation of the RI and FL tomograms. All the tomograms were manually inspected after the registration to be included in the dataset. It is worth noting that this procedure is not necessary after training and evaluation.

To reconstruct the stacked BF intensity images corresponding to the RI tomograms, we took advantage of the full optical field information measured by multi-angle interferometry (Mertz, 2019; Park et al., 2018). We first mapped the fields to a 2D Fourier space corresponding to the focal plane, thereby obtaining an amplitude and phase image with suppressed coherent noise due to the synthetic aperture. This 2D field was numerically refocused by convolving the propagation kernel with a varying distance of propagation. The BF intensity at each plane was calculated as the square of the amplitude image. We used MATLAB (MathWorks) to implement the BF reconstruction script.

### Model design, training, and inference

We used a single network architecture for all subcellular targets in order to avoid optimizing individual target-specific architectures. We automatically designed a highly flexible architecture through a scalable neural architecture search (Kim et al., 2019) (SCNAS). Specifically, SCNAS utilizes a stochastic sampling algorithm in a gradient-based bi-level optimization framework to jointly search for the optimal network parameters at multiple levels with generic 3D medical imaging datasets. As a result, a U-Net-like encoder-decoder structure with skip connections was discovered (Figure S1). At the end of every micro-level architecture, known as a motif, we added a dropout operation. The network parameters were as follows: activation function, leaky ReLU; normalization function, instance normalization; size of initial feature map, 12; number of layers, 8; feature map multiplier, 3.

To promote precise inference of both the large- and small-scale structures in the FL tomograms, for network training, we used a loss function, *l*, with both mean squared error (MSE), *l_MSE_*, and gradient difference loss (GDL), *l_GDL_*, terms: *l* = *l_MSE_* + *l_GDL_*. Each term is defined as follows:

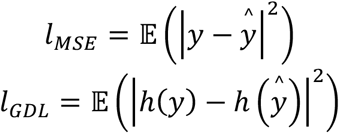

where *y* and 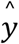 are the ground truth and inferred FL channel, respectively, *h*(·) is the 3D Sobel operator, and 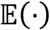 is the expectation over voxels and operations.

To train the networks, we used an Adam optimizer with an initial learning rate of 0.001, where the learning rate was reduced by a factor of 5 if there was no improvement in the validation metrics for 30 epochs. Randomly sampled parameters were used for data augmentation techniques such as flip, rotation, cropping, elastic deformation, and gamma correction. Hyperparameter optimization was based on a grid search algorithm whose search space consisted of hyperparameter combinations with similar memory and FLOPS requirements (Tan and Le, 2019). We used PyTorch in Python 3 to implement the deep learning pipeline.

Due to the memory constraints of GPU computing, we trained the networks using 3D patches instead of the whole tomograms. During training, the patches were randomly cropped from regions with registered FL data. For post-training inference, an RI tomogram was symmetrically padded, divided into overlapping patches with regular spacing, individually processed by the networks, and then stitched into whole FL tomograms with a spline kernel-based blending (Figure S2). The default size of a patch was 256×256×64 voxels.

### Performance and uncertainty quantification

Three performance metrics were used, namely peak signal-to-noise ratio (PSNR), Pearson correlation coefficient (PCC), and structural similarity index (SSIM). Each metric is defined as follows:

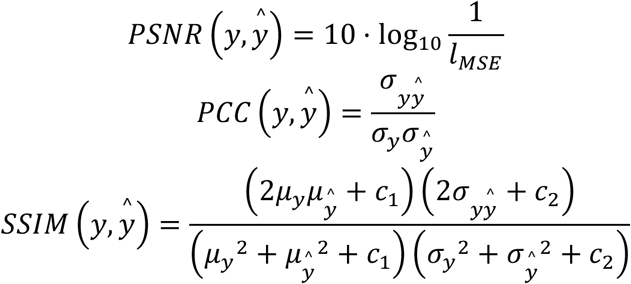

where *μ* and *σ* are the mean and standard deviation, respectively, covariance 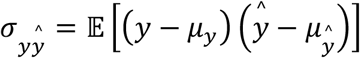, *c*_1_ = 0.01^2^, and *c*_2_ = 0.03^2^ by default. Following convention, we used a non-zero minimum standard deviation for PCC and a 3D Gaussian kernel with a size of 7 voxels for SSIM. The metrics were complementary to each other. PSNR is relevant for MSE, but suffers from poor perceptual performance and noise vulnerability. PCC has a decent perceptual performance, but hardly captures local differences. SSIM, which is relatively complicated, is a comprehensive metric designed to overcome the shortcomings of PSNR or PCC. SSIM can be factored into three terms called luminance, contrast, and structure. The luminance term is similar to MSE but uses mean values instead of voxel values. The contrast term quantifies the similarity of high-frequency components relevant to the GDL. The structure term is nearly identical to PCC. Various quantifications of the three metrics are shown in Table S2.

Following recent Bayesian deep learning approaches for computer vision, two types of uncertainty were considered, namely data (aleatoric) and model (epistemic) uncertainty. While the precise origins and mathematical derivations have been extensively reviewed elsewhere (Kendall and Gal, 2017), here, we describe the uncertainty quantification schemes well-suited for RI2FL. Data uncertainty was quantified by test-time augmentation (Wang et al., 2019) with image transforms, such as flip and rotation, which was compatible with the aforementioned loss function. Model uncertainty was quantified using a Monte Carlo dropout (Gal and Ghahramani, 2016). In both cases, we quantified the mean and standard deviation in the FL output space upon the perturbation of either the data or model. The two calculated standard deviation maps defined the data and model uncertainty. The average of the two mean prediction maps defined the final inferred FL, which slightly increased the performance as well. We did not apply these schemes to the stitching or time-series (except for the cell division example) data due to high computational costs.

### Single-cell profiling

Upon inference with RI2FL, a variety of open-source computational tools developed for FL data could be readily utilized. To segment the single cells and nuclei in the tomograms, we first trained a random forest voxel classifier, provided by Ilastik (Berg et al., 2019), based on the inferred nuclei and plasma membranes channels. The voxels were sparsely annotated as background, cytoplasm, or nucleus for a handful of tomograms, and the trained classifier generated the voxel-wise class probability maps for the entire dataset. Then the single nuclei could be readily segmented by thresholding the nuclei probability. The tentative cells, obtained by thresholding the summation of the cytoplasm and nuclei probability, were segmented by marker-controlled watershed segmentation, provided by CellProfiler (Bray et al., 2016), using the identified nuclei as segmentation markers. The segmentation performance was robust due to the high specificity of the inferred FL channels.

We extracted a variety of single-cell features from the segmented single cell/nuclei volume masks, the additional inferred FL channels (actin, mitochondria, lipid droplets, and nucleoli), and the measured RI channel aligned in a common coordinate system. The calculation of the mass-related features was based on the well-characterized linear dependence of RI, *n*(*x,y,z*), to the dry mass density, *C*(*x,y,z*), for biological samples (Barer, 1953; Park et al., 2018): *n*(*x,y,z*) = *n_m_* + *αC*(*x,y,z*), where *n_m_* and *α* are the RI of the imaging medium (*n_m_* = 1.337 at λ = 532 nm) and the RI increment (*α* = 0.190 mL/g at *λ* = 532 nm), respectively. The mass density and actin density described in the main text indicate the cellular dry mass density and cytoplasmic actin mean, respectively, in Table S3. For the time-lapse experiments, we used the framewise application of the segmentation and feature extraction procedures to quantify the feature dynamics. We used MATLAB (MathWorks) to implement the feature extraction script.

Uniform manifold approximation and projection (UMAP), implemented in Python 3, was used for the unsupervised nonlinear embedding of the features for 2D visualization (McInnes et al., 2018). All features were z-scored before UMAP, and the hyperparameters were as follows: minimum distance, 0.5; number of neighbors, 5.

### Data and code availability

Data and materials availability: RI2FL source code and data are available at https://github.com/NySunShine/ri2fl.

---

**Figure S1.**
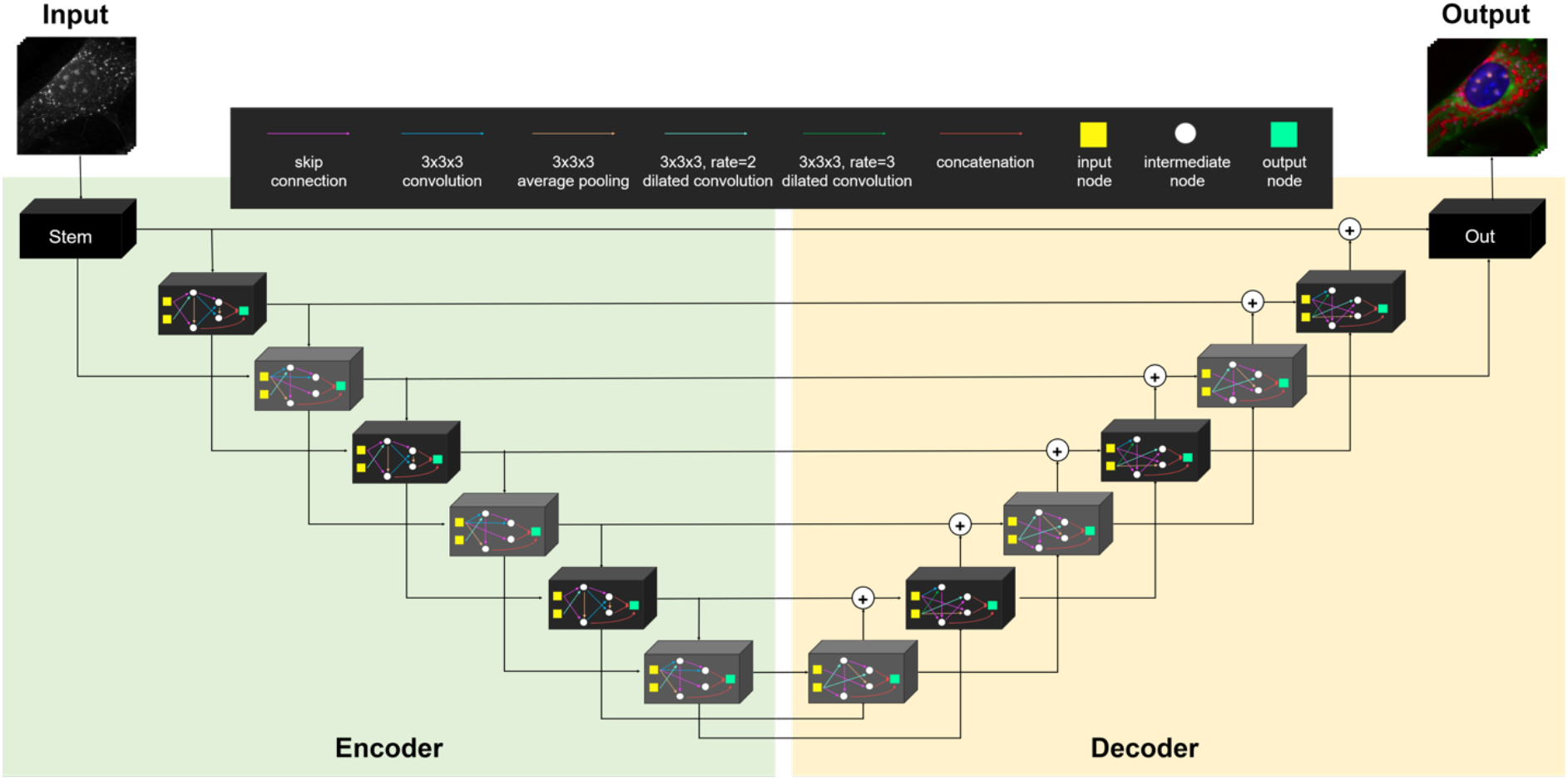
Network architecture. We used a single encoder-decoder network, systematically discovered by SCNAS, for all subcellular targets.

**Figure S2.**
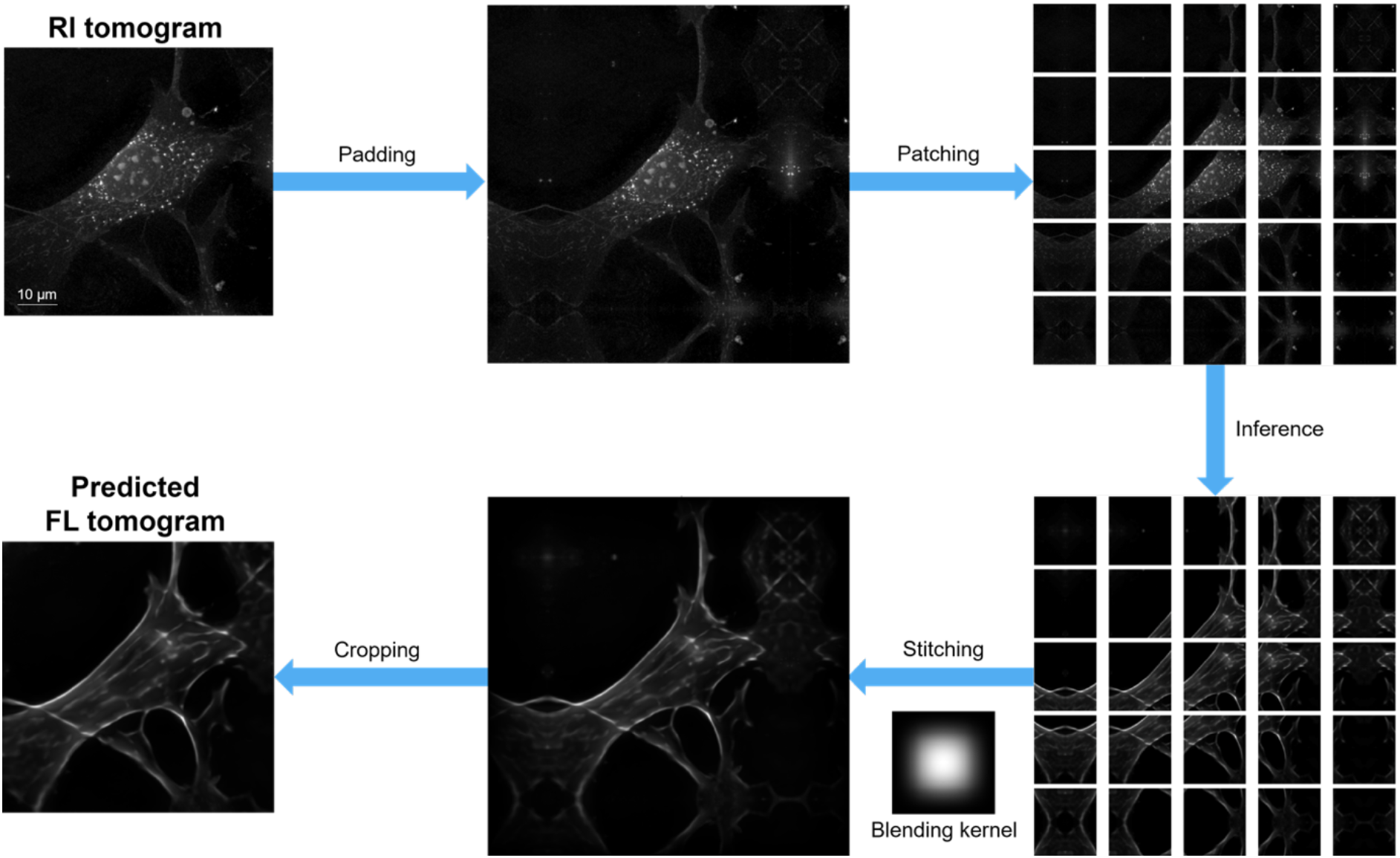
Patch-based processing. An example inference from an RI tomogram to the corresponding actin tomogram is illustrated as a flow chart, highlighting the patch-based processing for GPU memory management. All images represent maximum intensity projections of 3D data.

**Figure S3.**
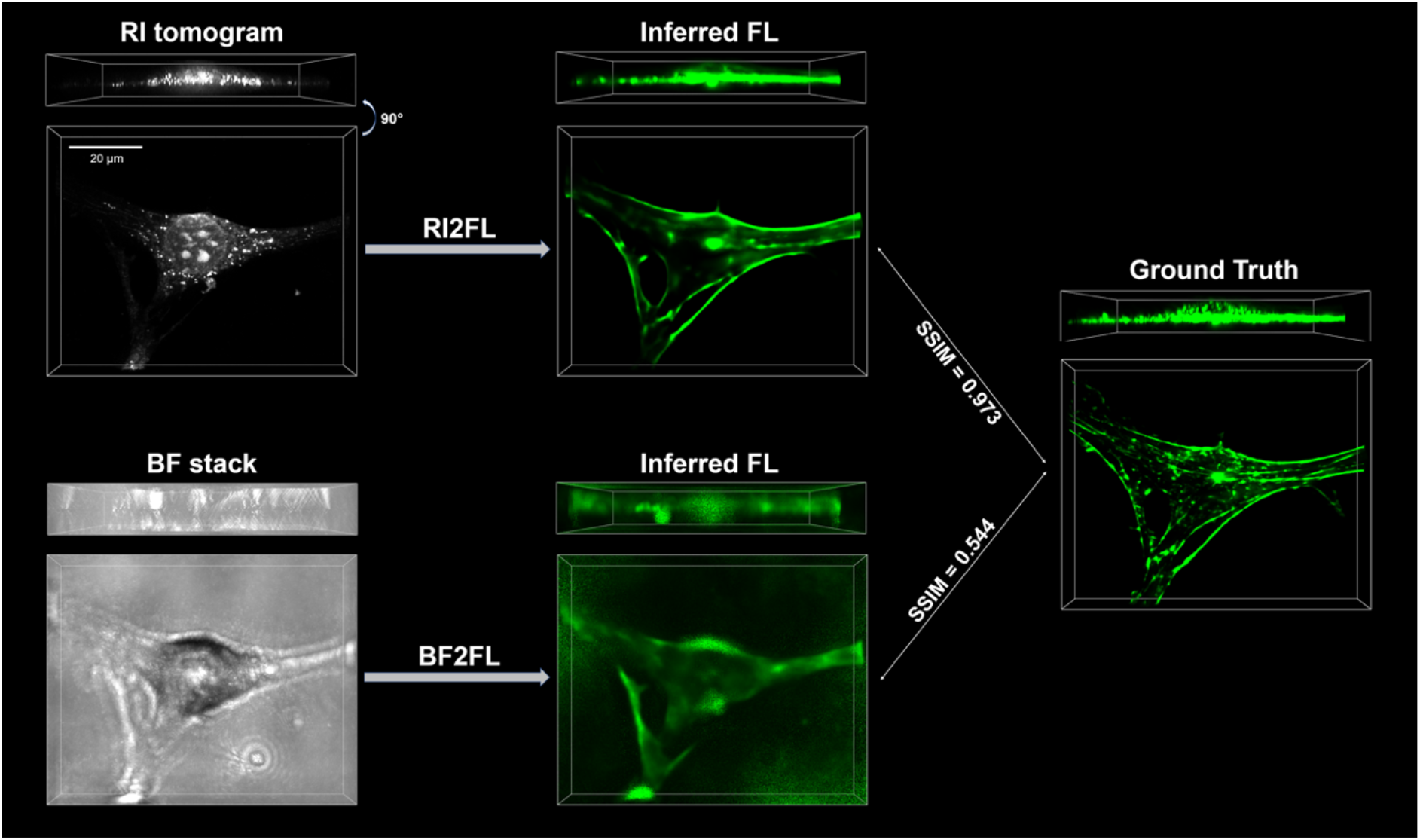
Comparison between RI2FL and BF2FL. An example inference with ground truth actin tomogram is shown. Clearly, RI tomograms enable more accurate FL inference compared to BF stacks, as quantified in Figure 2C. The bounding boxes represent 76.8×62.1 ×12.8 μm^3^ volumes corresponding to an identical FOV.

**Figure S4.**
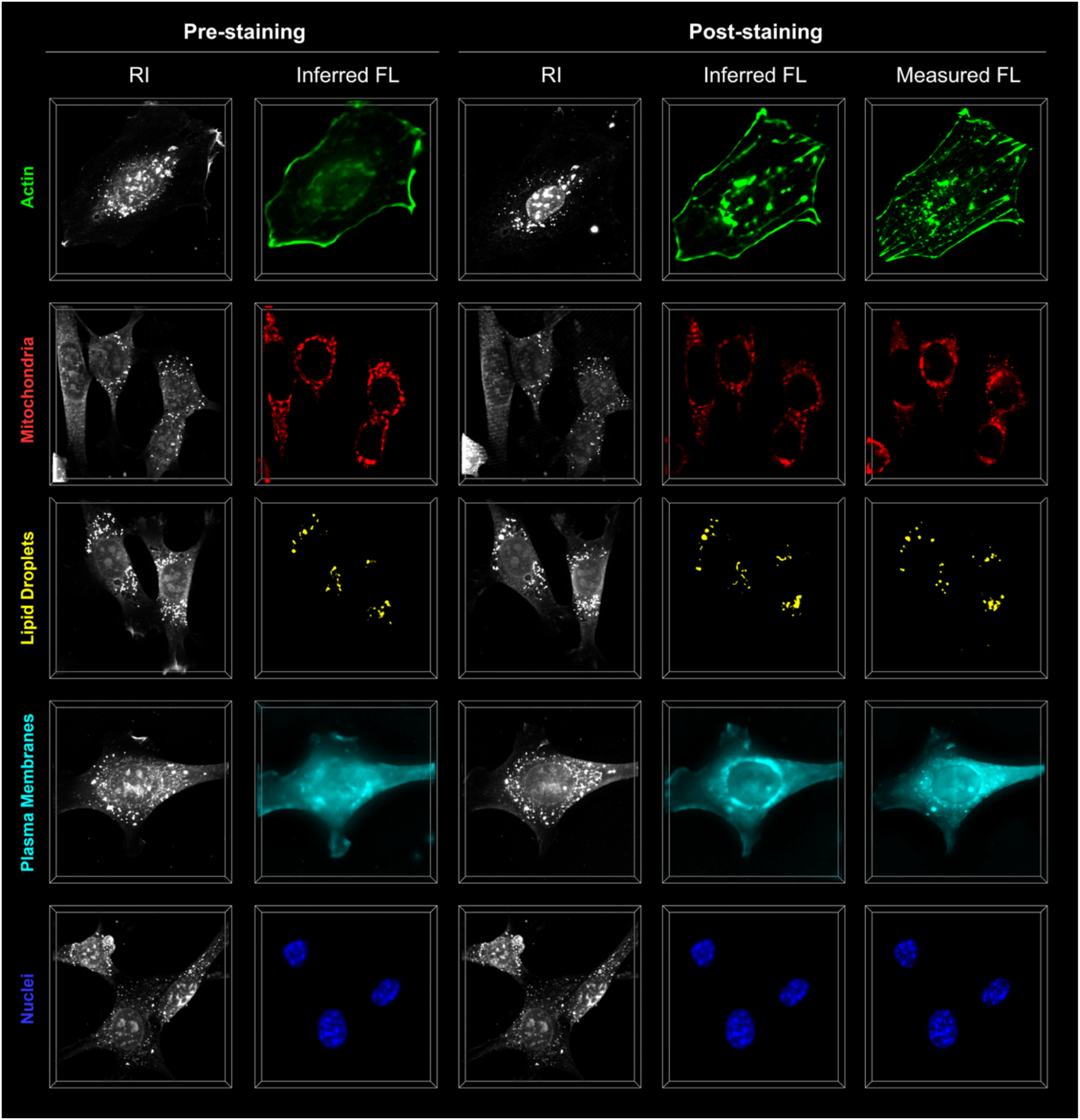
Comparison between pre- and post-staining data. In order to validate the inference of endogenous subcellular targets, we imaged identical cells before and after staining. The qualitative correspondence between the pre- and post-staining data, despite the irreducible discrepancy due to the temporal difference (from minutes to an hour) and fixation, further validates the successful operation of RI2FL in unlabeled cells.

**Figure S5.**
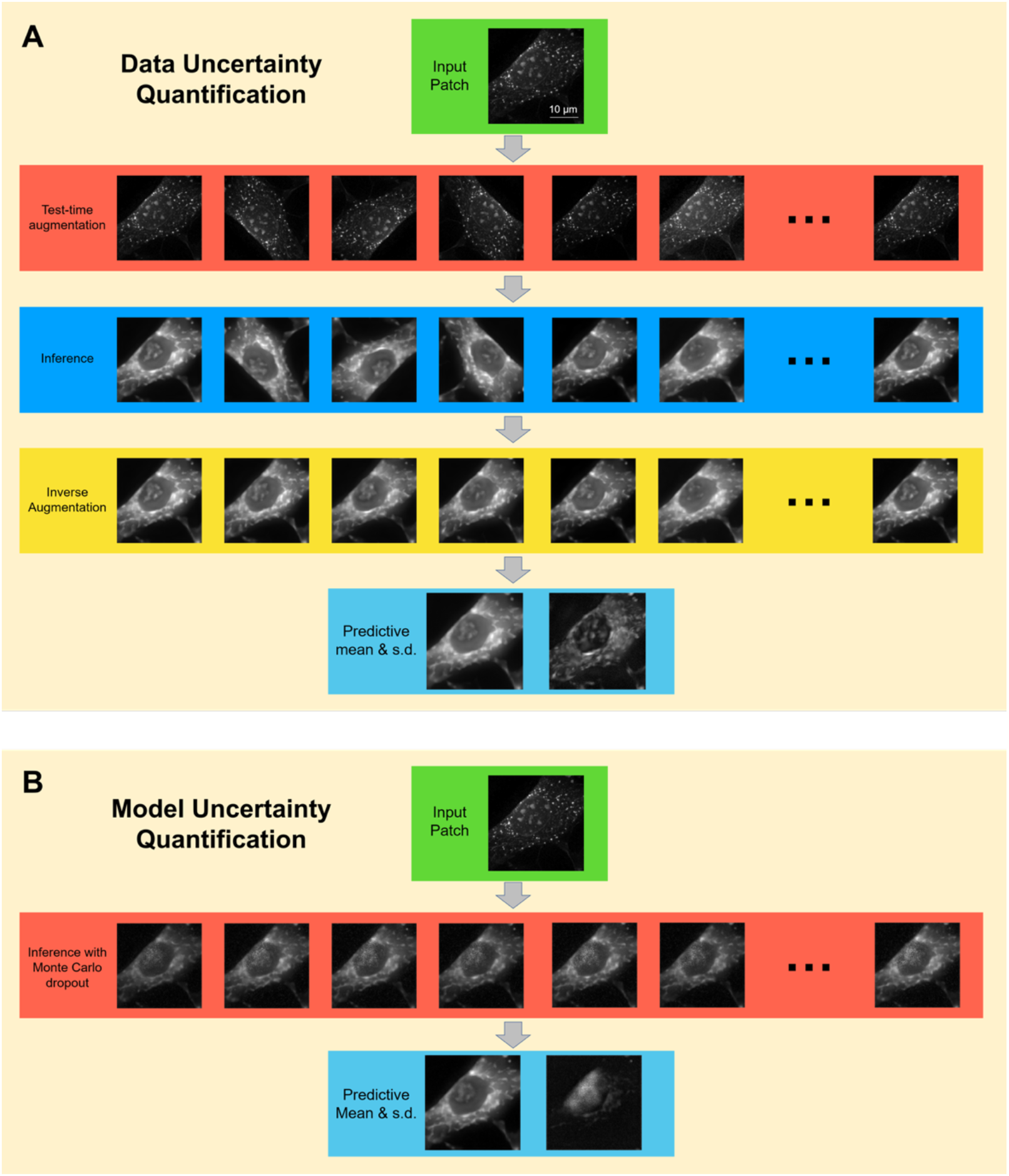
Uncertainty quantification schemes. (A) Data uncertainty and (B) model uncertainty quantifications were conducted by test-time augmentation and Monte Carlo dropout, respectively. All images represent maximum intensity projections of 3D data.

**Figure S6.**
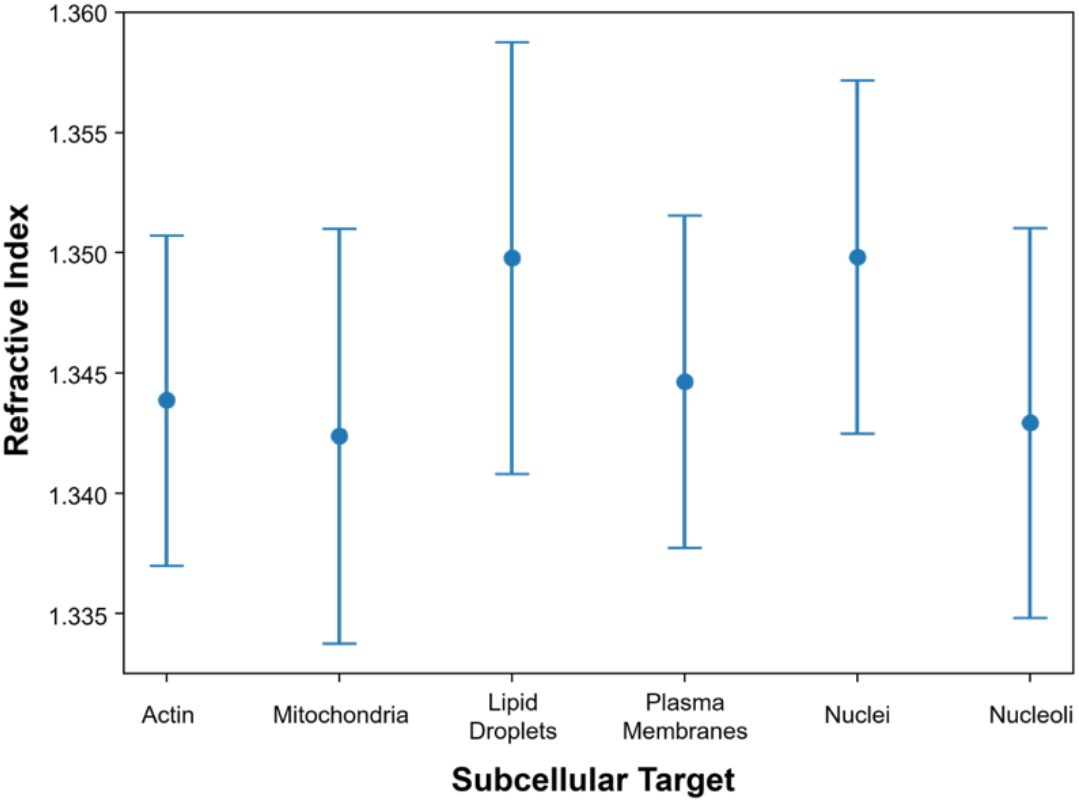
RI ranges of the subcellular targets. The target-specific RI ranges were estimated using the inferred FL data. For each FL channel, the voxels corresponding to the targets were determined by Otsu’s method. Mean±S.D.; the results were calculated from 703 NIH3T3 tomograms.

**Figure S7.**
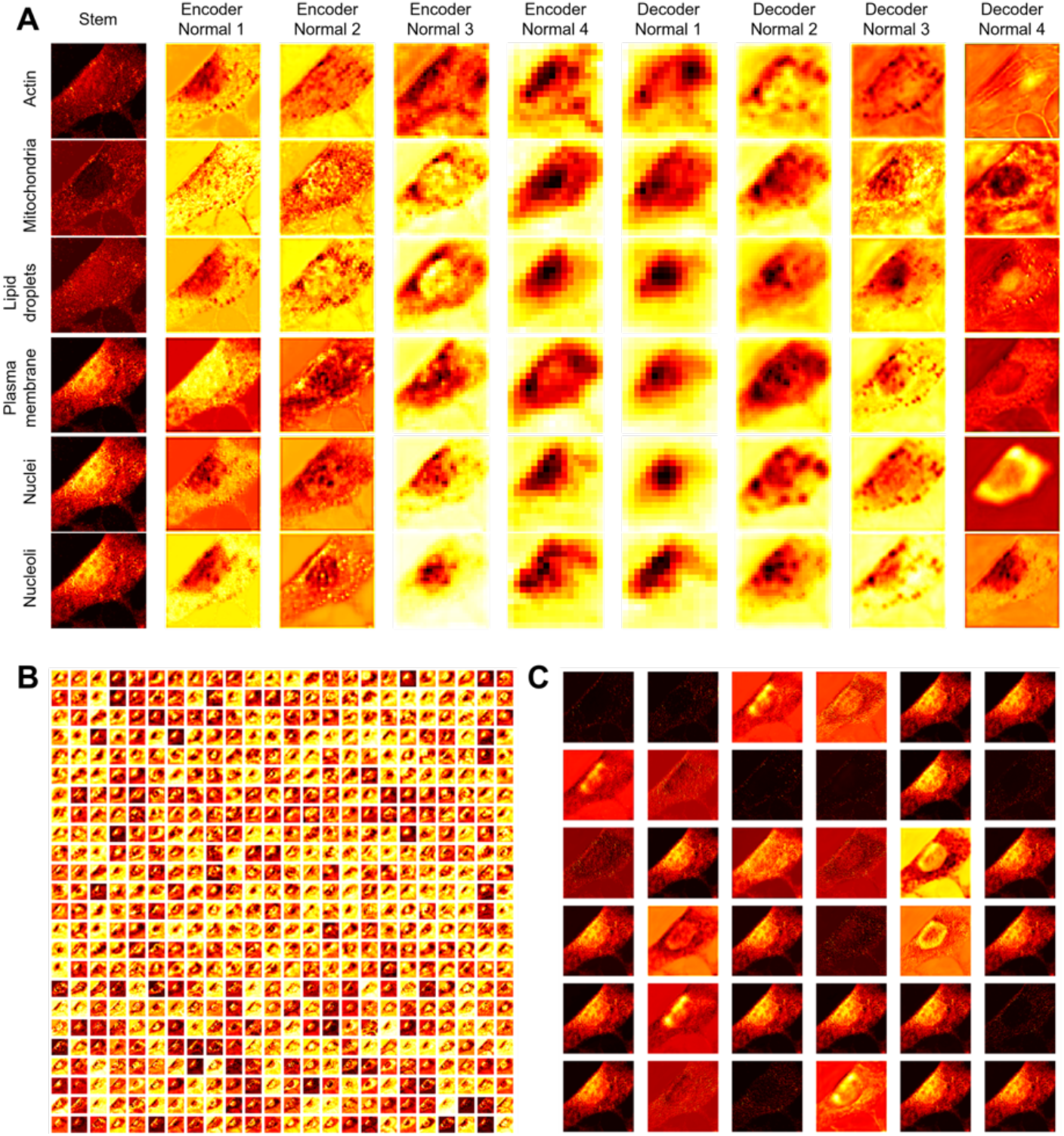
Deep feature visualization. In order to facilitate interpreting the operation of the trained networks, feature map activations for a single input tomogram were visualized. (A) Average feature map at each layer. (B and C) Individual feature maps at the last layers of encoder (B) and decoder (C) parts of the network inferring plasma membranes. All images represent maximum intensity projections of 3D data.

**Table S1.** Dataset summary.

| (Number of Tomograms) |  | Actin | Mitochondria | Lipid Droplets | Plasma Membranes | Nuclei | Nucleoli | Total |
| --- | --- | --- | --- | --- | --- | --- | --- | --- |
| Training | NIH3T3 | 98 | 248 | 84 | 116 | 83 | 69 | 698 |
| Validation | NIH3T3 | 10 | 10 | 7 | 10 | 5 | 5 | 47 |
| Test | NIH3T3 | 11 | 10 | 7 | 10 | 5 | 5 | 48 |
|  | COS-7 | 65 | 104 | - | - | 79 | - | 248 |
|  | HEK 293 | 3 | 55 | - | - | 50 |  | 108 |
|  | HeLa | 112 | 86 | - | - | 20 | 45 | 263 |
|  | MDA-MB-231 | - | 62 | - | - | 87 | - | 149 |
|  | Astrocyte | - | 14 | - | - | 26 | - | 40 |
| Total |  | 299 | 589 | 98 | 136 | 355 | 124 | 1,601 |

**Table S2.** Performance metrics.

| (SSIM, PSNR, PCC) |  | Actin | Mitochondria | Lipid Droplets | Plasma Membranes | Nuclei | Nucleoli | Average |
| --- | --- | --- | --- | --- | --- | --- | --- | --- |
| BF2FL | NIH3T3 | **0.541 ± 0.040** | **0.854 ± 0.043** | **0.865 ± 0.021** | **0.449 ± 0.101** | **0.537 ± 0.059** | **0.810 ± 0.143** | **0.635 ± 0.186** |
|  |  | 27.1 ± 0.9 | 30.0 ± 1.2 | 28.8 ± 0.7 | 23.3 ± 1.0 | 24.3 ± 0.9 | 30.7 ± 1.4 | 26.9 ± 2.9 |
|  |  | 0.269 ± 0.113 | 0.185 ± 0.030 | 0.093 ± 0.042 | 0.252 ± 0.105 | 0.362 ± 0.189 | 0.184 ± 0.115 | 0.229 ± 0.130 |
| RI2FL | NIH3T3 | **0.935 ± 0.019** | **0.951 ± 0.020** | **0.988 ± 0.005** | **0.924 ± 0.034** | **0.986 ± 0.005** | **0.918 ± 0.091** | **0.947 ± 0.042** |
|  |  | 35.9 ± 1.8 | 36.2 ± 2.2 | 43.8 ± 3.4 | 34.3 ± 2.2 | 38.7 ± 3.0 | 37.7 ± 3.9 | 37.3 ± 3.9 |
|  |  | 0.804 ± 0.078 | 0.734 ± 0.046 | 0.893 ± 0.030 | 0.936 ± 0.049 | 0.958 ± 0.028 | 0.755 ± 0.126 | 0.841 ± 0.105 |
|  | COS-7 | **0.811 ± 0.048** | **0.867 ± 0.031** | - | - | **0.878 ± 0.066** | - | **0.856 ± 0.056** |
|  |  | 28.1 ± 3.8 | 32.2 ± 2.4 | - | - | 26.6 ± 2.9 | - | 29.4 ± 3.9 |
|  |  | 0.448 ± 0.079 | 0.447 ± 0.060 | - | - | 0.693 ± 0.163 | - | 0.525 ± 0.157 |
|  | HEK 293 | **0.831 ± 0.033** | **0.755 ± 0.067** | - | - | **0.775 ± 0.109** | - | **0.766 ± 0.089** |
|  |  | 28.7 ± 1.3 | 27.6 ± 2.6 | - | - | 23.2 ± 2.6 | - | 25.6 ± 3.4 |
|  |  | 0.525 ± 0.064 | 0.384 ± 0.119 | - | - | 0.611 ± 0.155 | - | 0.493 ± 0.176 |
|  | HeLa | **0.821 ± 0.066** | **0.840 ± 0.039** | - | - | **0.820 ± 0.056** | **0.870 ± 0.063** | **0.836 ± 0.059** |
|  |  | 28.3 ± 2.7 | 29.7 ± 2.1 | - | - | 24.8 ± 1.8 | 36.8 ± 2.0 | 29.9 ± 4.1 |
|  |  | 0.473 ± 0.139 | 0.462 ± 0.066 | - | - | 0.538 ± 0.117 | 0.506 ± 0.218 | 0.480 ± 0.138 |
|  | MDA-MB-231 | - | **0.808 ± 0.049** | - | - | **0.889 ± 0.031** | - | **0.855 ± 0.056** |
|  |  | - | 29.7 ± 3.4 | - | - | 26.3 ± 1.8 | - | 27.7 ± 3.1 |
|  |  | - | 0.412 ± 0.088 | - | - | 0.635 ± 0.065 | - | 0.542 ± 0.133 |
|  | Astrocyte | - | **0.776 ± 0.032** | - | - | **0.940 ± 0.026** | - | **0.883 ± 0.084** |
|  |  | - | 31.4 ± 2.7 | - | - | 29.3 ± 4.6 | - | 30.1 ± 4.1 |
|  |  | - | 0.406 ± 0.056 | - | - | 0.613 ± 0.141 | - | 0.541 ± 0.155 |
|  | Average | **0.824 ± 0.064** | **0.829 ± 0.063** | **0.988 ± 0.005** | **0.924 ± 0.034** | **0.866 ± 0.082** | **0.875 ± 0.066** | **0.845 ± 0.074** |
|  |  | 28.7 ± 3.5 | 30.4 ± 3.2 | 43.8 ± 3.4 | 34.3 ± 2.2 | 26.2 ± 3.6 | 36.9 ± 2.2 | 29.3 ± 4.5 |
|  |  | 0.484 ± 0.142 | 0.441 ± 0.098 | 0.893 ± 0.030 | 0.936 ± 0.049 | 0.644 ± 0.142 | 0.531 ± 0.223 | 0.529 ± 0.167 |

(Mean±S.D.; the numbers of tomograms are shown in Table S1)

**Table S3.**
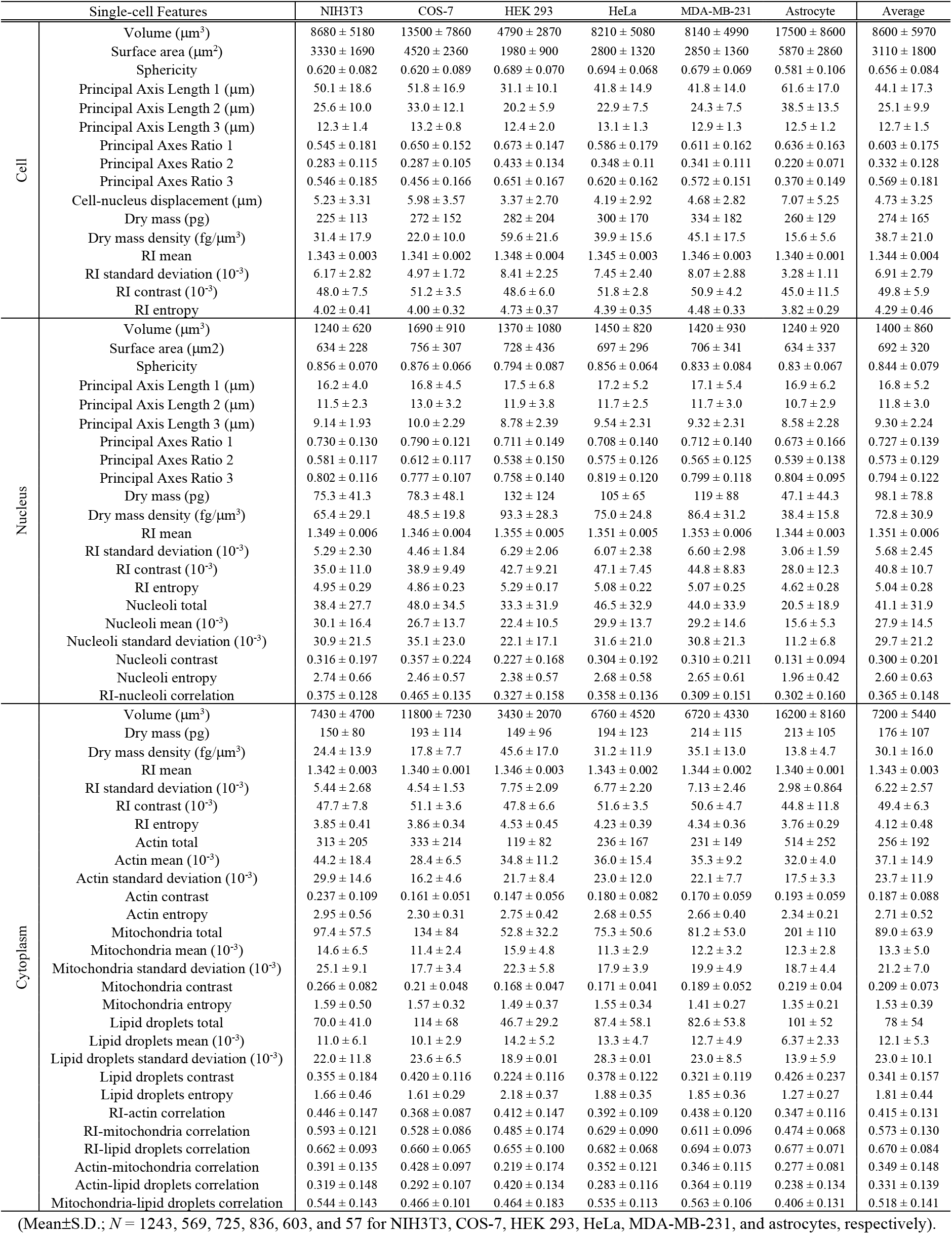
Single-cell feature statistics.

## Video Captions

**Video S1.** RI2FL across cell types and subcellular targets.

**Video S2.** Dynamics of cell division and RI2FL.

**Video S3.** Large-scale imaging and RI2FL.

**Video S4.** Growth factor stimulation.

**Video S5.** Chemogenetic stimulation.

